# Force generation by polymerizing filaments revisited: diffusive interaction leads to nonlinear force-number scaling

**DOI:** 10.1101/125690

**Authors:** Jemseena Valiyakath, Manoj Gopalakrishnan

**Affiliations:** Department of Physics, Indian Institute of Technology, Madras, India

**Keywords:** polymerization, filament, stall force, microtubule, diffusive interaction

## Abstract

Polymers growing against a barrier generate force and push it forward. We study here force generation by a bundle of *N* rigid polymers growing in parallel against a diffusing, rigid, flat barrier, resembling a bundle of microtubules. To estimate the polymerization force, the barrier is subjected to a force *f* acting against the direction of growth of the polymers and the mean velocity *V_N_* (*f*) of the filament assembly is computed. The maximum polymerization force (*alias* stall force) 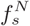 is deduced from the condition 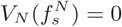. This problem has been studied in the literature earlier, but two important aspects have escaped attention: (a) free diffusion of monomers is hindered by the barrier, even when it is far from the growing tips and (b) parallel filaments could interact through the monomer density field (“diffusive coupling”), leading to negative interference between them. In our model, both these effects are investigated in detail. A mathematical treatment based on a set of continuum Fokker-Planck equations for combined filament-wall dynamics suggests that the barrier reduces the influx of monomers to the growing polymer tip, thereby reducing the growth velocity and also the stall force, but it doesn’t affect the scaling of the stall force with number, i.e., 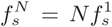. However, Brownian dynamics simulations show that the linear scaling holds only when the filaments are far apart; when they are arranged close together, forming a bundle, sublinear scaling of force with number appears. We argue that the nonlinear scaling could be attributed to diffusive interaction between the growing tips which becomes significant when the tips are close together. These conclusions, initially established for simple flat-faced polymers, are also found to hold true for microtubules with their characteristic hollow cylindrical geometry and rugged tip structure. In particular, simulations show conclusively that the stall force of a single microtubule is a fraction of the combined stall force of the 13 protofilaments. This result is supported by a simple analytical estimate of the force using diffusive coupling theory, and is in agreement with earlier experimental observations.

## 1 Introduction

The cytoskeleton, a network of filaments composed of actins and microtubules forms one type of force generators inside a eukaryotic cell and is crucial in moving cyotoplasmic material within the cell [1]. One manifestation of such force generation includes formation of the mitotic spindle, an apparatus formed by microtubules with associated molecular motors during cell division where the genetic material (chromosome pairs) are being pulled and pushed by microtubules until the sister chromatids are separated to each daughter nuclei [2, 3]. Yet another instance is the migration of a cell from one place to another by crawling; in this case, polymerizing actin filaments pushing the plasma membrane of the cell provides the mechanism for motility[4, 5].

Outside the cytoplasmic environment, even a minimal system comprising of biofilaments growing against a barrier (rigid or elastic) has been shown to be capable of generating forces of the order of a few piconewtons [6–11]. For instance, a single microtubule grown *in vitro* can generate a maximum of 5pN force against a rigid barrier [6]. Microtubules growing against obstacles, tend to bend from their straight trajectory and then proceed to grow; a condition known as buckling [6, 8]. Mechanically induced changes stemming from confinement are observed to alter the intrinsic dynamics of microtubules [8, 12]. The catastrophe transition becomes pronounced in the vicinity of the barrier [12, 13], while the dynamic instability is observed to be regulated by force [8].

It is well known that the physical barrier (the plasma membrane or proteinous kinetochore *in vivo*) in contact with the microtubule will affect the filament dynamics, by rendering steric hindrance to further polymerization. Evidence for this scenario may be found in the reduced growth velocity near the cell boundary, as reported by [13] in fission yeast cells; here, the reduction in growth velocity is speculated to be mechanical in origin. However, a slightly different explanation could be offered. Prior to the assembly, the monomers are diffusing in the cytoplasm or the available space of the experimental chamber, and any hindrance to free diffusion would have observable consequences; in particular, growth rate may be reduced due to hindrance to free diffusion offered by the barrier. This scenario appears to be partly supported by observations which find that growth of microtubule in the interior of the cell is different from near the boundary [14]. This effect is distinct from the steric hindrance to monomer addition, which comes into play only at extremely small barrier-tip separation (of the order of monomer length). A clear distinction between these two cases is not apparent from the existing experimental observations, and therefore, we investigate it in detail here.

The origin of the polymerization force is to be traced to the free energy change associated with polymerization[1]. For example, the free energy released per GTP-tubulin addition to a microtubule is nearly 5 – 10 k_B_T, equivalent to a force of 50 pN if a microtubule grows by 8nm; similarly the gain in free energy per GDP-tubulin dissociation from a microtubule is nearly 5 – 10 k_B_T [11]. However, experimental measurements suggest that the force produced by a microtubule is only a fraction of the maximum force predicted by theoretical arguments, suggesting that not all the free energy available from polymerization of the 13 protofilaments is converted to work[6]. Obtaining insights into this interesting nonlinear scaling behaviour is another important motivation for us in undertaking the present study.

With the primary objectives having been outlined above, it would be appropriate to summarize our methods and important results here, before going into the details of both: we first study a mathematical model for collective dynamics of a bundle of filaments that grow via diffusion limited adsorption in a confined space, against a mobile barrier that is subjected to an external force acting against the direction of growth of the polymers, as well as thermal noise. Unlike previous models, we do not treat the growth rate of the filament as a constant parameter; rather, it is regarded as a continuous function of the distance from the barrier, derived by solving the steady state diffusion problem in the presence of absorbing (filament tip) and reflecting (wall) surfaces. A number of analytical results are derived from this model, built on continuum Fokker-Planck equations for combined barrier-filament dynamics. In particular, for monomers in the shape of flat-faced disks (FFD), the model predicts that barrier-induced hindrance is significant if the radius of cross section of the disk *a* ≫ *δ*, the length of the monomer. In this case, both growth velocity and force of the filament assembly is reduced by this effect; but the combined force still scales linearly with the number of filaments. Support to these predictions, as well as further insights are provided by Brownian dynamics simulations, which are performed first with FFD filaments (with various radii of cross-section), and later with filaments with microtubule-like geometry. The simulations clearly show that the force-number scaling depends on the lateral separation between the filaments; when the filaments are placed closed together, the scaling is sublinear. We argue that the origins of this sublinear scaling of force with number may be traced to a subtle effect, i.e., *diffusive interaction* between the filament tips, which arises out their mutual competition for monomers. Specifically, we find that a simple theoretical expression based on diffusive coupling successfully predicts the reduced stall force of a single microtubule (when compared with the combined force of the 13 protofilaments).

## 2 Materials and Methods

### 2.1 Model details

Our model consists of a bundle of *N* filaments growing from one fixed wall of a compartment, towards the opposite wall, which is a diffusing barrier (diffusion coefficient *D_ω_*), also being pushed backward with a constant force *f*, see Fig.1. The filaments grow by diffusion-limited polymerization of monomers, which are present in the solution at concentration *C_0_*. A filament also shrinks by random detachment of monomers, with rate *k*_off_. The filaments in the bundle are identical, and have equal base separation from each other. No interaction is assumed to exist between the filaments or between a filament and the barrier, except that neither of them can penetrate each other (see [15–17] which explicitly considers the energy of interaction between the filaments). The filaments are assumed rigid, unaffected by thermal noise. We do not include the chemical switching activity of the monomers, similar to some earlier studies [16, 18–20], our filaments are chemically inert.

**Figure 1:**
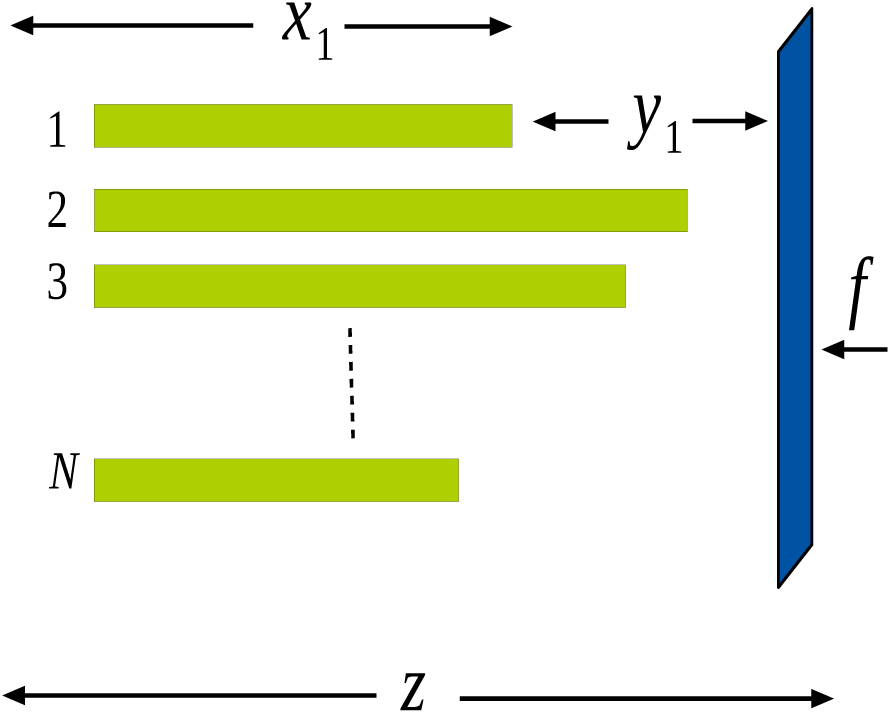
Schematic diagram for a bundle of inflexible filaments pushing against a movable rigid barrier acted upon by a constant force *f*. The rigid barrier also undergoes thermal motion characterised by diffusion coefficient *D_ω_*.

### 2.2 Mathematical formalism: Fokker-Planck equations

Most of the mathematical models that dealt with polymerization-driven force generation do not explicitly consider the wall movements. Rather, the presence of wall is encapsulated in the growth rate and detachment rate of the filament [16, 17, 19–21]. In these ‘Brownian ratchet’ models, the filament in contact with the wall is assumed to grow with an on-rate proportional to exp (–*qfδ*/*k_B_T*) and off-rate proportional to exp (–(*q* – 1)*fδ*/*k_B_T*) with ‘*q*’ being the load sharing factor. Here, our approach is different; we adopt a formalism similar to [18]. In this model, the wall executes a combination of diffusive (arising from thermal noise) and directed (due to the external force) motion. Also, a continuum approximation is adopted for the filament dynamics as we find it is convenient to incorporate the continuous variation of on-rate with the wall-filament separation (discussed in more detail later).

The joint probability density function *P*(*X, z*; *t*) for the filament tip positions *X* ≡ {*x*_1_, *x*_2_, .., *x_N_*} and wall position *z*, in the continuum limit, satisfies the diffusion-drift equation

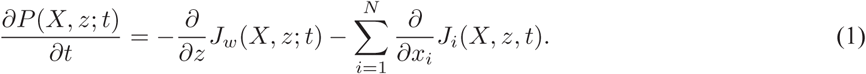

Eq.1 is a Fokker-Planck equation in *N* + 1 variables, with both *x_i_* and *z* lying in the interval (−∞,+∞). The individual probability currents corresponding to the dynamics of wall and filaments are given by

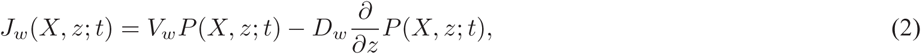

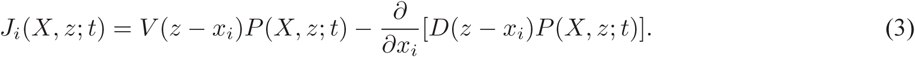

The ‘diffusion coefficient’ *D* and ‘velocity’ *V* are expressed in terms of the on- and off-rates of monomers by the standard expressions [22]

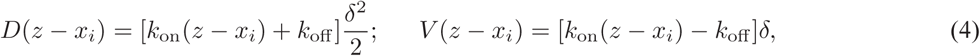
where *k*_on_(*z* – *x_i_*) is the position-dependent on-rate for monomer adsorption to the filament tip, the position dependence arising from the modification of the steady state concentration field due to the presence of the barrier (see Appendix A for details). *k*_off_, the off-rate of monomers is assumed to be a constant. Using fluctuation-dissipation theorem, the wall velocity *V_ω_* = –*fD_ω_*/*k_B_T*. The currents are subjected to the boundary conditions

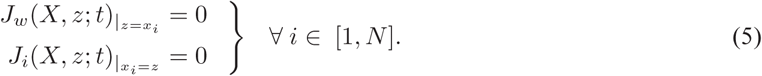

Let the instantaneous separation between the *i^th^* filament and the wall be denoted

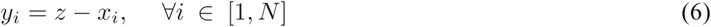
which we shall refer to as ‘gaps’. Given the reflecting boundary conditions in Eq.5, we expect the gap probability distribution (see Eq.15 later) to become stationary for non-zero *f*, in the long-time limit. This conjecture helps us derive an expression for our main quantity of interest, i.e., the average filament/wall velocity, in a straightforward way. Considering this, we implement a change of variables in Eq.1. All *x_i_* are thus transformed into *y_i_* by the relations given by Eq.6. We denote the transformed probability density function as Π(*Υ, z*; *t*), where *Y* = {*y*_1_, *y*_2_, ., *y_N_*}, hence Eq.1 becomes,

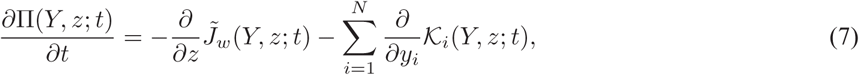
where, 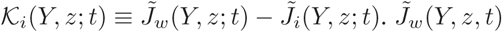 and 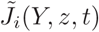 are the transformed probability currents, in terms of the new variables, which are, respectively,

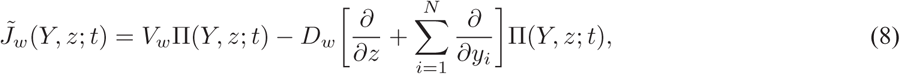

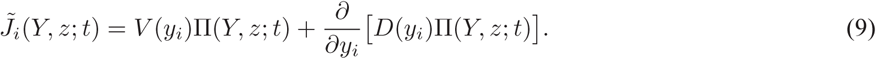
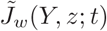 and *K_i_*(*Y*, *z*; *t*) satisfy the boundary conditions (for 1 ≤ *i* ≤ *N*)

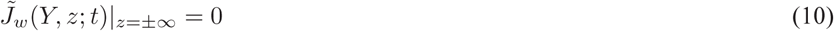

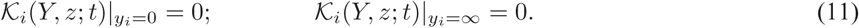

The average position of the wall is given by

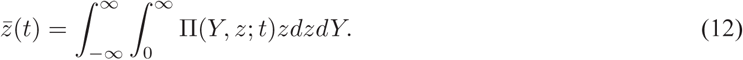

Applying Eq.12 in Eq.7 and using the boundary condition given by Eq.10 and 11, we arrive at the following expression for the steady state mean wall velocity

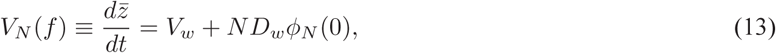
where *ϕ_N_*(*y*) is the stationary probability density for the separation *y* between the wall and one of the filaments (single filament gap size distribution), i.e.

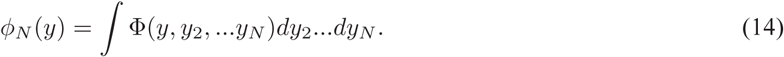

The integrand Φ(*Y*; *t*), gives the joint probability distribution of gap lengths:

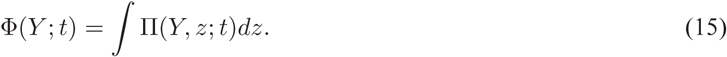

From Eq.6, it follows that in the long time limit, the average wall velocity becomes equal to the average filament velocity, hence it is sufficient to get an expression for the wall velocity using Eq.13, using which one can study the wall-induced effects on the kinetics of polymerization and force generation.

### 2.3 Brownian dynamics simulations

The mathematical formalism presented earlier has two limitations: (i) diffusion of monomers is not taken into account explicitly, rather, it enters through the gap-dependent on-rate of monomers (ii) the length of the polymers is treated as a continuous variable, ignoring the discreteness of monomer addition and dissociation processes. To overcome these limitations, we also carried out Brownian dynamics simulations; here, the free monomers are treated as point particles, and diffuse inside a rectangular box, with the walls of the box acting as reflecting boundaries, see Fig.2.

**Figure 2:**
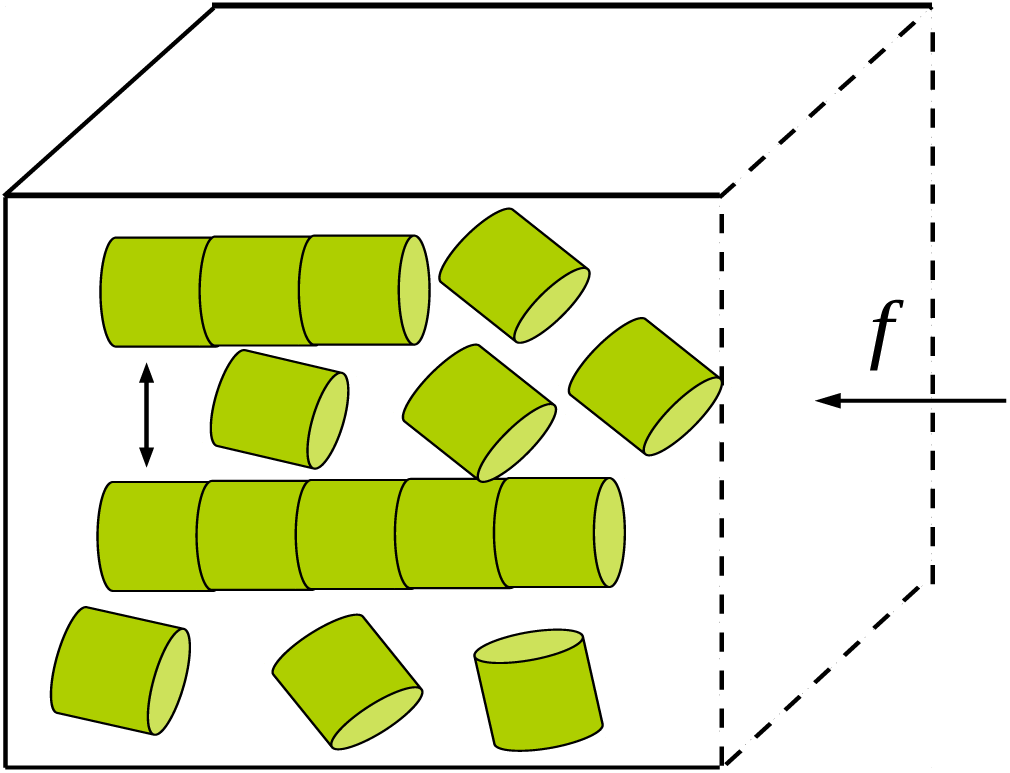
Schematic diagram of the cubical box, containing a bundle of filaments growing by a diffusion-limited reaction used in Brownian dynamics simulations. One face of the cubical box facing the filament tip is movable (the barrier); it is acted on by a constant force *f* in the backward direction and also undergoes random motion characterised by diffusion coefficient *D_ω_*.

The positions of the individual monomers **r***_m_*(*t*) and the wall (movable face of the rectangular box) are updated using overdamped Langevin equations. Over a small time step Δ*t*, the updating rules have the form,

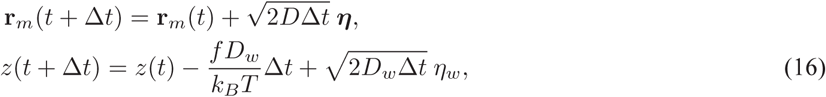
where ***η*** = (*η_x_, η_y_, η_z_*), the latter being random numbers drawn from independent Gaussian distributions with zero mean and unit variance. Similarly, *η_ω_* is a Gaussian random variable with zero mean and unit variance. We used Δ*t* = 10^−4^ s, *D_ω_* = 10^3^ nm^2^s^−1^ and *D* = 10^5^ nm^2^s^−1^; for diffusing monomers, this implies a mean free path 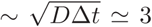 between successive changes in direction. A ratio of 100 : 1 was kept between the diffusion coefficients of the wall and the monomers since the objects that obstruct the free growth of microtubule as well as free diffusion of tubulins include the lipid membranes or a kinetochore, which are massive compared to tubulin monomers. Three boundary conditions are imposed on the diffusing monomers, (i) reflecting boundary conditions on the walls of the rectangular box, (ii) reflecting boundary condition on the cylindrical wall of the filament and (iii) absorbing boundary condition at the circular face/tip of the filament. Although the monomers are treated as point particles when simulating their diffusion, once a monomer is adsorbed onto a polymer tip, the length of the polymer increases by *δ* (however, this requirement was waived in one set of simulations, see (i) below). The initial spatial distribution of free monomers is uniform, with concentration *C_0_*. In order to ensure that adsorption events at polymer tips do not cause depletion in the *total* number of monomers, every time a monomer disappeared from the solution by binding to a polymer tip, a new monomer was added at a random location inside the box. This procedure ensures that the free monomer concentration far from the absorbing tips is always *C_0_*. In the simulations, we analyzed three different cases and are summarized below.

#### (i) On-rate of monomers binding to static disk-shaped absorbing surface

In the first set of simulations, we looked at the steady state adsorption of particles to a static filament (filament length remains the same irrespective of monomer adsorption) with all the faces of the box kept fixed, by varying the separation between the filament tip and the face opposite to it. Here, the filament as well as the wall are static in space.

#### (ii) FFD filaments growing against a mobile barrier

In the second set of simulations, we studied the force-velocity relation for rigid linear polymers with monomeric units modeled as flat-faced disks of different radii of cross section. In addition, we also varied the base separation between the filaments. Unlike the earlier case, here, whenever a monomer is adsorbed, the length of the filament increases by *δ*; the mean length of a filament grows linearly with time. We also allow a bound monomer to occasionally detach from the filament after adsorption, and this is accounted for in the simulations by including a non-zero off-rate. Our simulations are done at fixed *mean* monomer concentration *C_0_* (as explained earlier). The mean velocity of growth of a filament was measured as the slope of the graph of the mean length versus time, with the averaging done over 1000 independent runs. For different values of *f*, we calculated the average velocity of growth of the filament, for *a* = 20 nm, 10 nm and 2 nm, with *δ* = 2 nm in all the three cases. The mean velocity was plotted as a function of the force *f*; the stall force *f_s_*, the point of zero-crossing of the *V* – *f* curve, was determined by linear interpolation.

#### (iii) Multi-stranded polymers with microtubule-like geometry

In the next stage, we extended our simulations to multi-stranded polymers with microtubule-like geometry (but without hydrolysis or dynamic instability, see the schematic figure, Fig.3). Here, each polymer consists of 13 protofilaments (with each protofilament being a FFD polymer with radius *a* = 2.5 nm, similar to one of the cases studied in (ii)), arranged in a circular fashion, with outer radius 12.5 nm and inner radius 7.5 nm. Each protofilament here grows and shrinks individually, with diffusion-limited binding and random detachment (off-rate *k*_off_) of monomers. The monomers here are circular disks of radius 2.5 nm and length *δ* = 8 nm. To calculate the mean velocity of growth, we tracked the time evolution of the length of one randomly chosen protofilament belonging to one of the microtubules (if there are more than one) in one simulation. The results are averaged over 500 independent runs.

**Figure 3:**
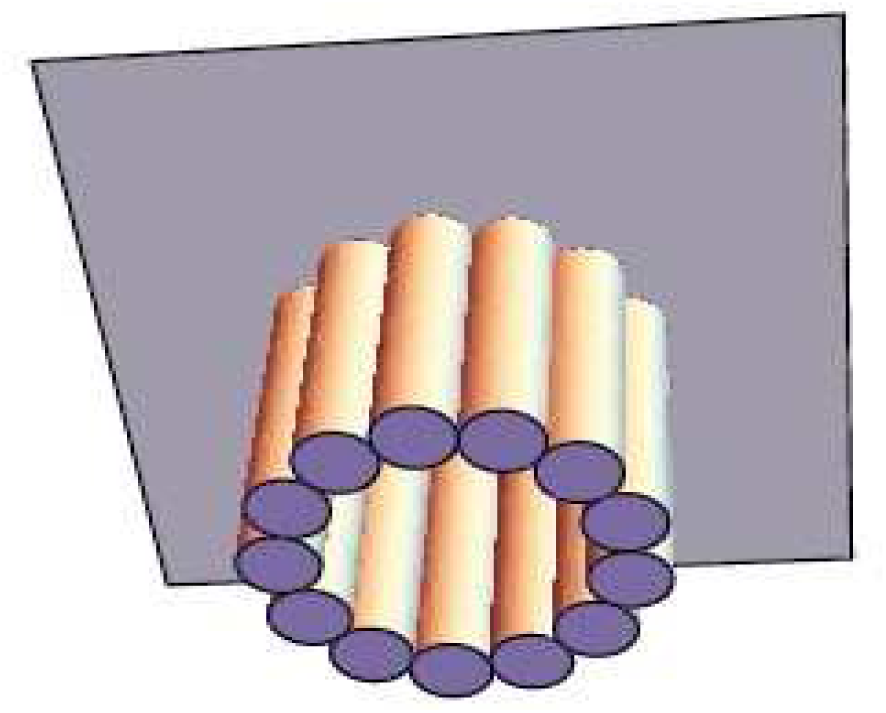
A schematic diagram of a multi-stranded filament with microtubule-like geometry.

## 3 Results

### 3.1 Mathematical results: Gap distribution and mean filament velocity

To derive an expression for the mean velocity of the filament/wall, we first need to find the expression for the single filament gap distribution, *ϕ_N_*(*y*). To derive the equation for *ϕ_N_*(*y*), we first integrate out *z* from the general equation for ∏(*Y*, *z*; *t*) given by Eq.7 and using the boundary conditions for 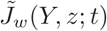 and 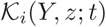 given by Eq.10-11 we get,

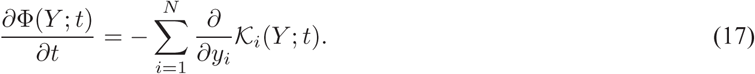

In the long-time limit, the gap size distribution Φ(*Y*; *t*) is expected to become stationary, hence the L.H.S of Eq.17 can be put to zero to give

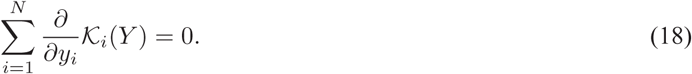

The simplest (but not the most general) solution to the above equation is 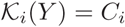, with *C_i_* being arbitrary constants. In this case, the boundary conditions given by Eq. 11 requires that *C_i_* = 0 identically. Hence, let us continue on the presumption that the set of equations

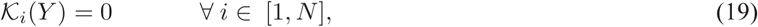
will provide the steady state solution for Φ(*Y*) we are looking for. Applying Eq.14 in Eq.19, we get the equation for single filament gap size distribution *ϕ_N_*(*y*) as

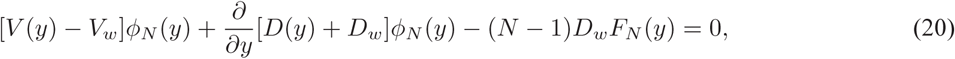
Where

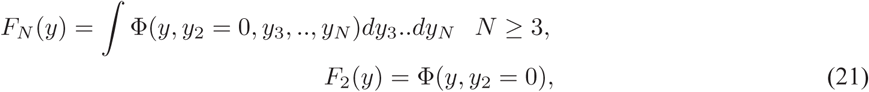
such that

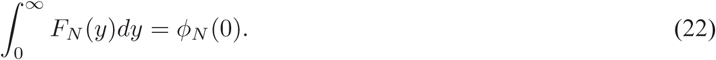

The formal solution to Eq.20 can be written as

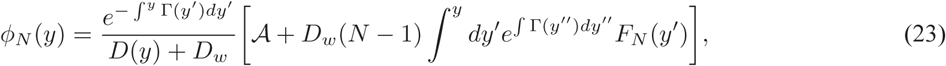
with

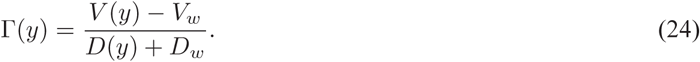

In order to calculate *ϕ_N_*(*y*), we use the following mathematical forms for *V*(*y*) and *D*(*y*), obtained using the approximate position-dependent on-rate *k*_on_ ≃ *k*_on_(∞)[1 – *e*^−λ*y*^] (see Appendix A) for FFD polymers. Here, we expect from scaling considerations that λ ∼ 1/*a*, where *a* is the radius of the disk. It then follows from Eq.4 that

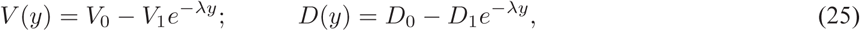
Where

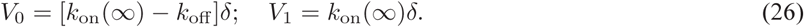

Similarly

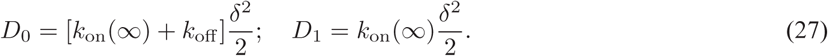

For *N* = 1, the gap size distribution is given by

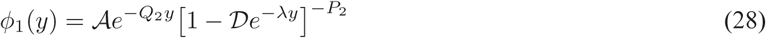
where

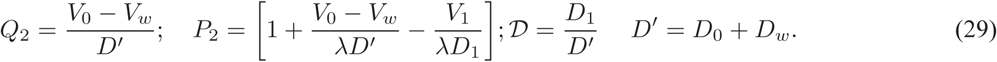

The normalization constant 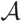 is given by

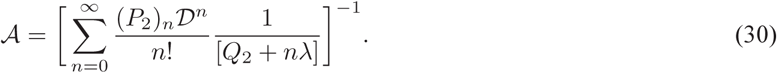

The corresponding results for *N* = 2 are given in *supplementary material*.

### 3.2 General solution for λ = ∞(constant on-rate)

The simplest limit, corresponding to λ = ∞, is similar to the studies by [18]; here, we have

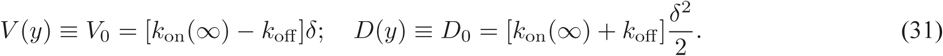

In this limit, the single filament gap distribution (Eq.23) is given by

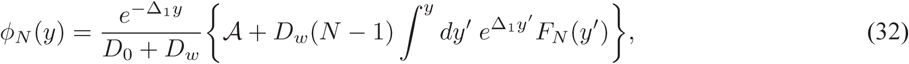
with

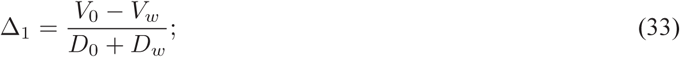

The expression for *F_N_* (*y*) is a simple exponential here:

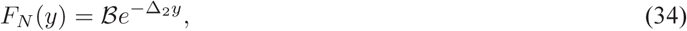
Where

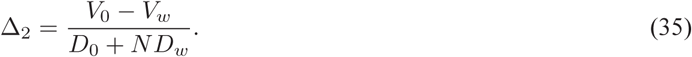

Now, substituting Eq.32 and Eq.34 in Eq.S16, we get

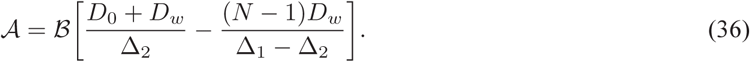

On using the normalization condition (Eq.S15) in Eq.32 we find that

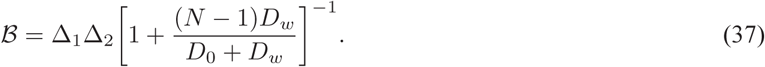

Substituting Eq.34 with 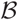 given by Eq.37 in Eq.32 and performing the integration we get

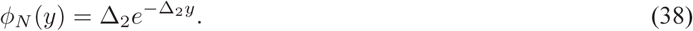

Substitution of *ϕ_N_*(0) = Δ_2_, as calculated using Eq.38, in the general expression for the average velocity given by Eq.13 gives

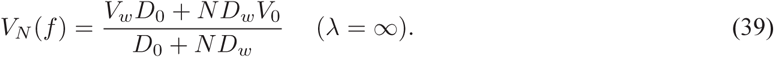

The stall force, corresponding to zero mean velocity, is obtained by putting *V_N_* (*f*) = 0 in Eq.39, and is given as

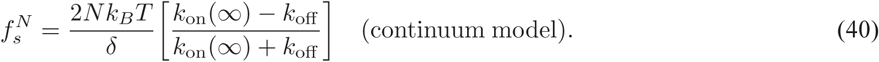

For constant on-rate case, the stall force scales linearly with the number of filaments, similar to earlier prediction [20]. But the mathematical dependence of stall force on the on-rate and off-rate differ, the difference clearly arising from the continuum treatment in this paper as opposed to the discrete approach in [20]. The corresponding prediction of the discrete model [20] is

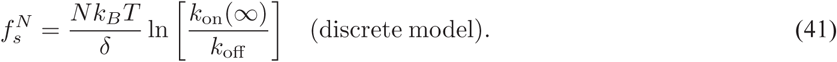

Note that *f_s_*(*N*) = 0 in both Eq.40 and Eq.41 when *k_on_* = *k_off_*. The latter corresponds to the state of chemical equilibrium of the system, where growth and detachment processes balance each other on average (with corresponding drop in the free monomer concentration in solution), and there is no net growth for the polymer (hence no more work can be extracted). It is also easily verified that in the limit *k*_on_(∞) ≈ *k*_off_, Eq.41 agrees with Eq.40, which is to be expected, as a continuum approximation works best when the (length) increment per unit step is small.

In Appendix B, we show that the linear scaling of stall force with number holds true to 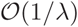, although both the filament velocity and the stall force are found to be reduced. These predictions are subjected to further examination in the following subsection.

### 3.3 Simulation results

In the first set of simulations we studied the on-rate of monomer adsorption to a static surface, in the presence of a reflecting wall, as mentioned in case (i) of Sec.2.3, for various radii of cross-section of the circular absorbing surface. From the simulation results, it is observed that the on-rate of monomers is dependent on the separation between the surface and the wall; as the separation decreases, a substantial drop in the on-rate is seen. The data along with the analytical results is discussed in detail in Appendix A, and was used in analytical calculations in the previous subsection.

In the second set of simulations, we studied the force-velocity relation for FFD filaments. Fig.4a shows the data for *a* = 10 nm, for one and two filament systems. In the second case, two values for the lateral base separation between filaments was studied; 0 and 100 nm (here, zero base separation refers to the filaments touching each other). The two cases are distinguished in the plots as “near” and “far”. The stall force for a single filament is found to be ≃ 1.94 pN. By comparison, Eq.41 predicts a stall force of ≃ 2.62 pN. The discrepancy is almost certainly arising from the barrier-induced reduction in the on-rate, which is significant for *a* = 10 nm. The inset of the same figure shows the dimensionless on-rate for monomer binding onto an absorbing disk of the same radius, as a function of its distance from a reflecting barrier. Fitting the data to an exponential curve yields the parameter λ, which is then used to predict the force-velocity relation using the relevant equations from Sec.3.1, viz., Eq.13, Eq.28 (for *N* =1) and Eq.S19 (*supplementary material*, for *N* = 2). The theoretical predictions are also shown alongside the simulation results in the same figure. For *N* = 1, the agreement between the theoretical curve and simulation data is excellent in the sub-stall regime, while significant deviation is observed post-stall. For *N* = 2, the far-data shows reasonable agreement with the theoretical curve except at very small *f*, but the near-data is significantly different. More importantly, while the two-filament stall force 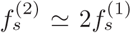 for filaments far apart, 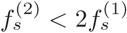 for near-filaments, i.e., sublinear scaling of stall force with number is observed when the filaments are close together, but linear scaling is restored when they are far apart. This observation is in disagreement with the prediction of the Brownian ratchet model [18]. We strongly believe that the sublinear scaling arises from *diffusive interaction* between the filaments, a phenomenon that occurs whenever multiple ‘sinks’ compete for diffusing particles that form a common pool [23–25].

**Figure 4:**
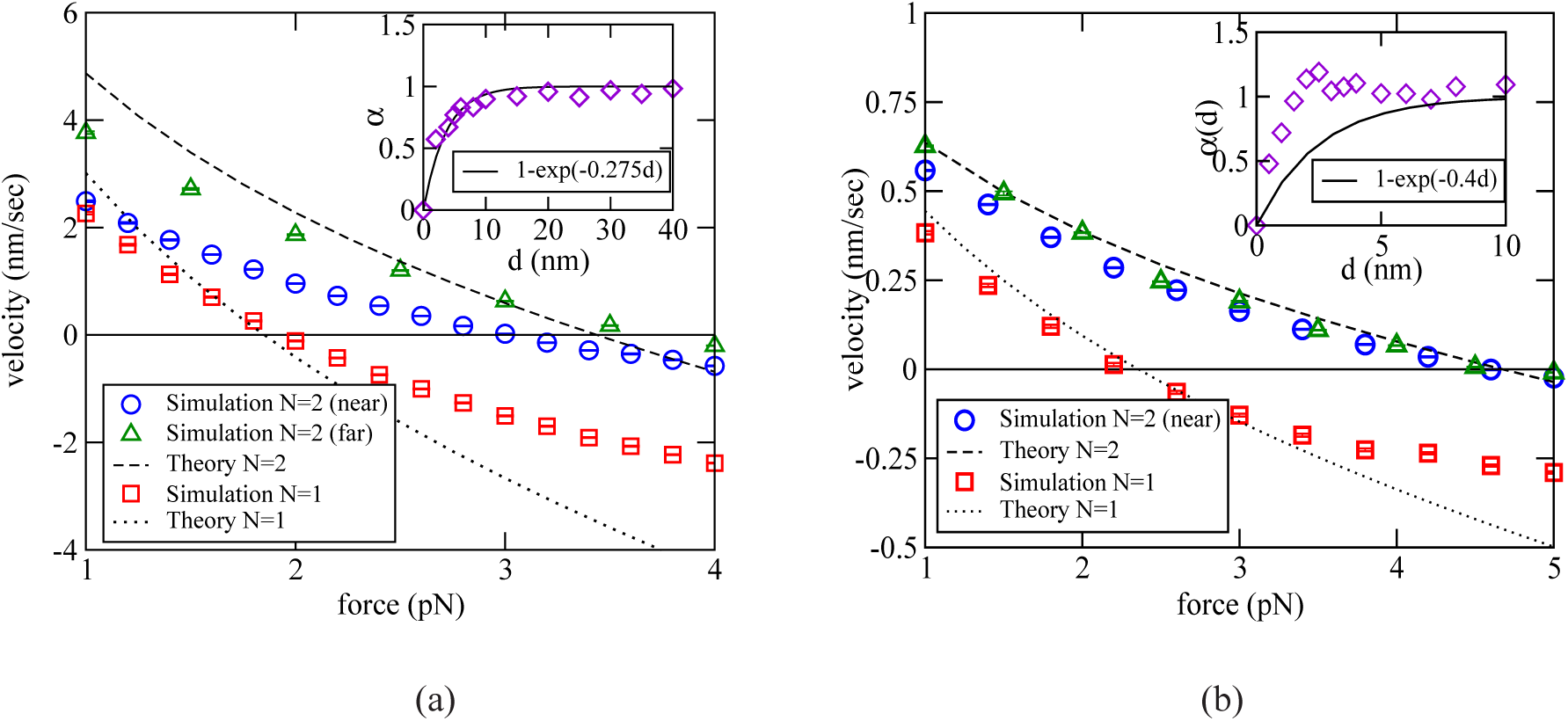
The force-velocity curve for a single filament versus two filaments, obtained from Brownian dynamics simulations, for (a) *a* = 10 nm and (b) *a* = 2 nm. For (a), the off-rate of monomers is *k*_off_ = 2 s^−1^ while for (b), *k*_off_ = 0.2 s^−1^. Analytical results are shown for best fit value of λ; 0.275 nm^−1^ in (a) and 0.4 nm^−1^ in (b). In the insets, fits for (scaled) separation-dependent on-rate *α*(*d*) = *k*_on_(*d*)/*k*_on_(∞), using the same λ are shown. The other parameter values are given in Table 1. Here, *near* means zero base separation between polymers, wheras *far* refers to a base separation 10*a*. The error bars are typically smaller than the size of the symbols.

We repeated the above investigations for a smaller radius of cross-section, *a* = 2 nm. The results are shown in Fig.4b. The observed single filament stall force here is nearly 2.27 pN, while the theoretical prediction from Eq.41 of the discrete model is ≃ 2.6 pN. The velocity-force curves predicted using the continuum model also agree with the simulations over a larger range of force. Coming now to two-filament data, unlike the previous case, the near and far cases for *N* = 2 are practically indistinguishable here, and both agree very well with the theoretical curve (here, “far” refers to a base separation of 20 nm). Finally, the two-filament stall force is very nearly twice the single filament force, indicating that diffusive interaction is negligible here, at least for *N* = 2. However, we shall see in the next subsection that for larger numbers, this interaction becomes significant even for *a* ∼ 2 nm. For two filaments, the observed doubling of stall force for *N* = 2, when the filaments are far apart, is also consistent with the results of the asymptotic analysis (λ → ∞) presented in Appendix B.

As further verification of the continuum theory presented in the last section, we also found the single-filament gap distribution function *ϕ_N_*(*y*) (defined in Eq.14), and compared with the theoretical predictions given in Eq.28 (*N* = 1) and Eq.S19 (*N* = 2). The results are given in Fig.5 (*a* = 10 nm) and Fig.6 (*a* = 2 nm). Quantitative agreement is better for *a* = 10 nm compared to *a* = 2 nm, as expected.

**Figure 5:**
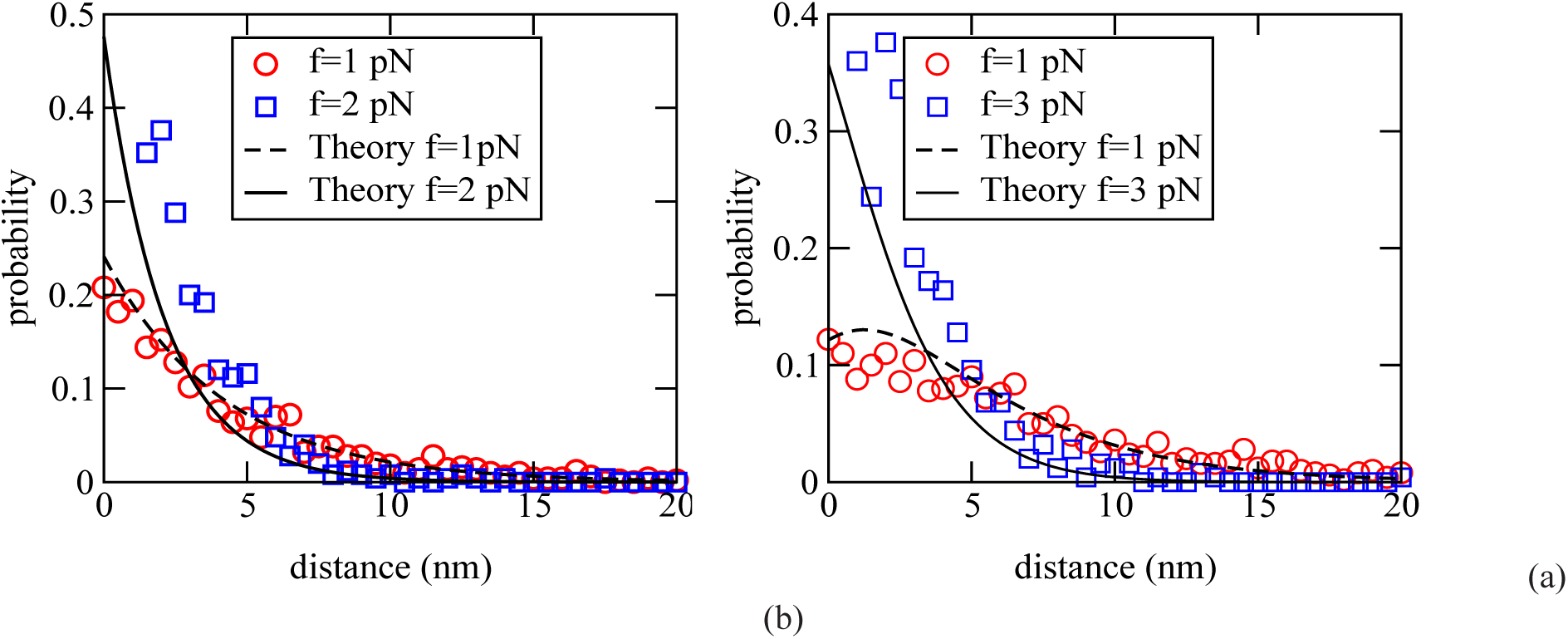
The gap distribution for *a* = 10 nm, for two forces, far from and near to stall, with (a) *N* =1 and (b) *N* = 2. Fits of the analytical results, Eq.28 for *N* = 1 and Eq.S19 for *N* = 2 are also shown for the best fit value λ = 0.275 nm^−1^, with *k*_on_(∞) = 7.5 s^−1^. For both (a) and (b), *k*_off_ = 2 s^−1^. The other parameter values are listed in Table 1.

**Figure 6:**
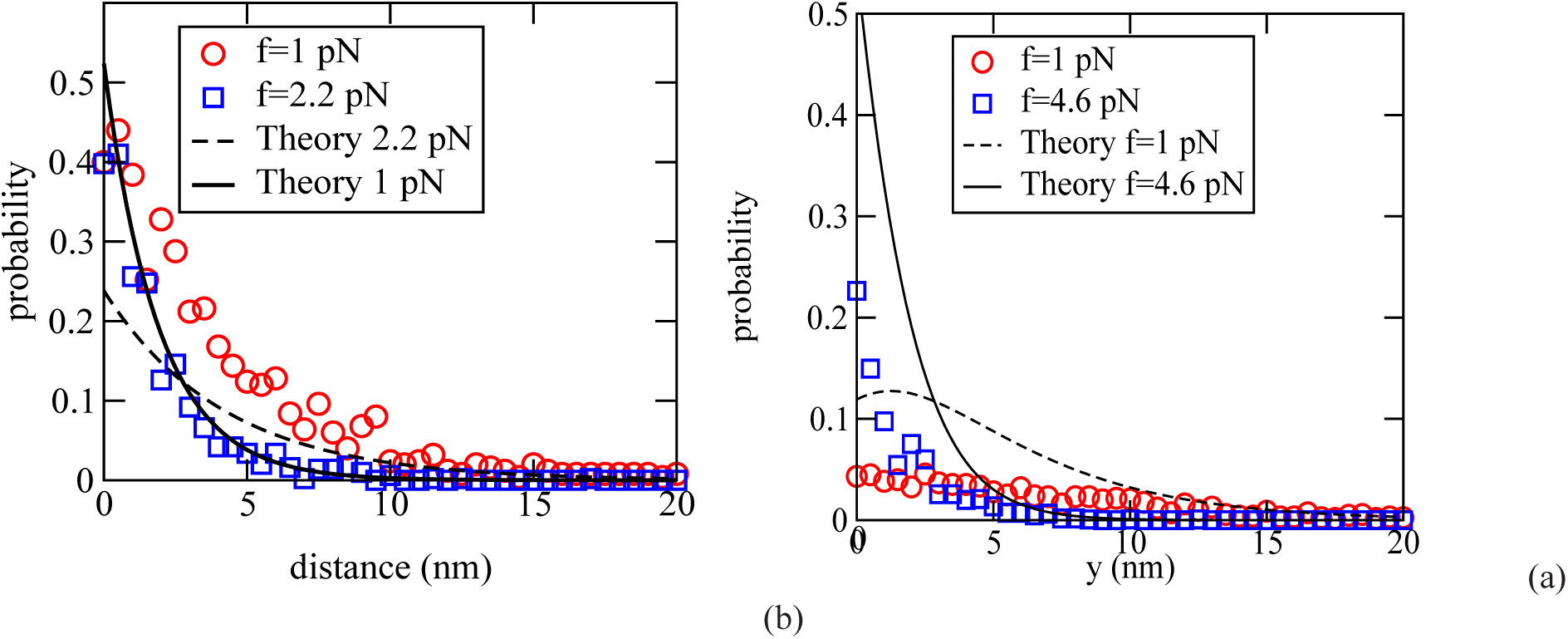
The gap distribution for *a* = 2 nm, for two forces, far and near to stall. (a) is shown for *N* = 1 and (b) is shown for *N* = 2. A fit of the analytical results (Eq.28 for *N* = 1 and Eq.S19 for *N* = 2) also shown for the best fitting value of λ = 0.4 nm^−1^, with *k*_on_ (∞) = 0.73 s^−1^. For both (a) and (b), *k*_off_ = 0.2 s^−1^. The other parameter values are listed in Table 1.

Fig.7 shows the force-velocity curve for one (*N* = 1) and two (*N* = 2) microtubule-like filaments. The observed stall force for a single microtubule is found to be nearly 5.17 pN in simulations for the parameters used here. For comparison, for the same set of parameters, the prediction of the Brownian ratchet model (Eq.41) for the combined stall force of 13 independent protofilaments is 12.24 pN. Therefore, we again encounter sublinear scaling of stall force, here as a function of the number of (proto)filaments, similar to the experimental observations in [6]. This can be explained using two different, but essentially equivalent arguments:

**Figure 7:**
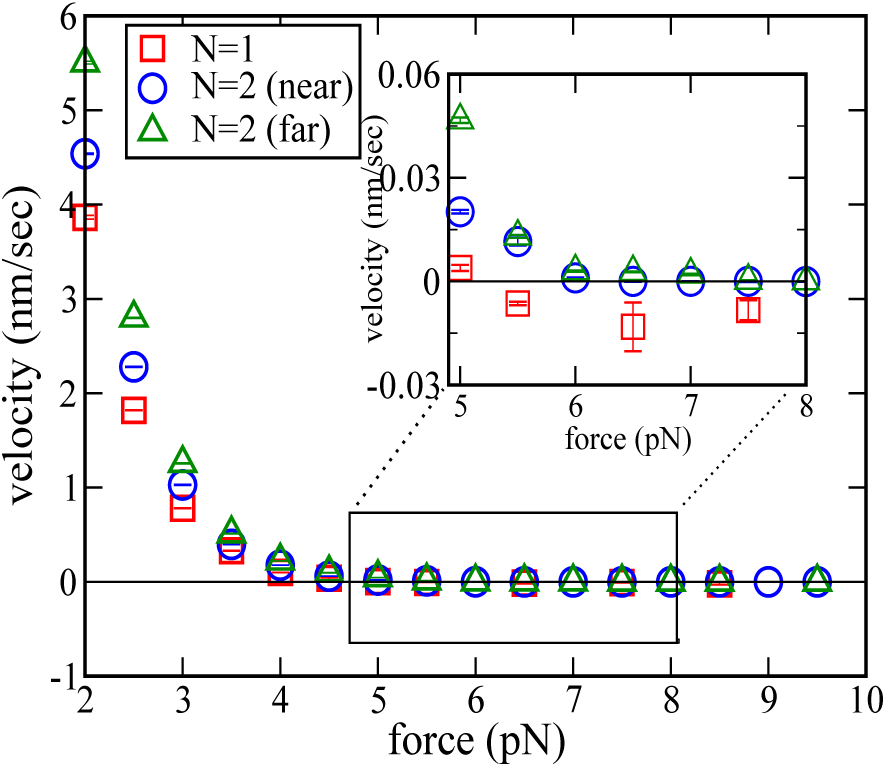
Force-velocity relation for a single microtubule and two microtubules, both *near* (zero base separation) and *far* (base separation of 150 nm). The values of the parameters used in the simulation are listed in Table 1. The inset zooms the force range where the velocity vanishes. Note that while the single filament curve crosses the *x*-axis after touching zero at stall, the two-filament velocity remains close to zero after stall. In most cases, the error bars are smaller than the size of the symbols. For two filaments, ‘near’ means zero base separation, while ‘far’ refers to a base separation 150 nm.

(a) a protofilament here is part of the larger microtubule, which has an outer radius of nearly 12.5 nm, large enough for significant barrier-induced reduction in the on-rate, when the filament is close enough to the wall. This causes each protofilament to grow much slower than it would have, if it were alone in the solution. Consequently growth is stalled at a lower value of the opposing force.

(b) each protofilament is diffusively coupled to the other protofilaments, and hence the on-rate for one is reduced by the presence of the others. For *n* disk-shaped absorbers (each with radius *a*) arranged uniformly in a circle, with centre-to-centre separation *R*, it has been proposed that, for *R* ≫ *a*, the effective diffusion-limited on-rate for one of the disks is given by the approximate formula [26]

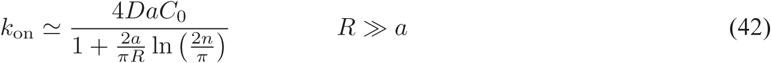

In a microtubule, protofilaments are tightly packed, this corresponds to the situation with *R* ≃ 2*a* in the above formula. However, protofilament lengths can be different in general, hence it is not clear *a priori* if Eq.42 can be applied in this case. Nevertheless, it is remarkable that the stall force of a single microtubule calculated using Eq.41, with the on-rate given by Eq.42 (after substituting *a* = 2.5 nm, *R* = 2*a, n* = 13 and *δ* = 8 nm), turns out to be 3.94 pN, closer to the observed value. The velocity-force curves of two microtubule-like filaments, both *near* (base separation zero) and *far* (base separation 150 nm), show a surprising feature. A close inspection (see inset of Fig.7) reveals that the two-filament mean velocity remains close to zero after reaching stall (|*V*_2_(*f*)| < 10^−3^ nm s^−1^ for 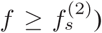. This counter-intuitive behaviour persists for forces up to 10 pN; the growth velocity remains close to zero for a large range of force in the super-stall regime. At present, we do not have an explanation for this observation. Nevertheless, Fig.8 provides some insights. Here, we show comparisons of the time-dependence of the mean position of the barrier and a randomly chosen protofilament for *N* =1 (a) and *N* = 2 (b and c), at *super-stall* forces. For *N* = 1, the wall and the filament keeps moves leftward on average, keeping a constant mean separation between them. Something different happens for *N* = 2. Here, as the force is increased, the mean positions of both the filament tip and the wall shifts leftwards, but settles in a new equilibrium position, with a constant, force-dependent mean separation between the two (Fig.8). Since both near and far configurations show similar qualitative behaviour, it appears that the large number of individual (proto)filaments for *N* = 2 (26 in total) might be the crucial factor here; this issue requires further investigation.

**Figure 8:**
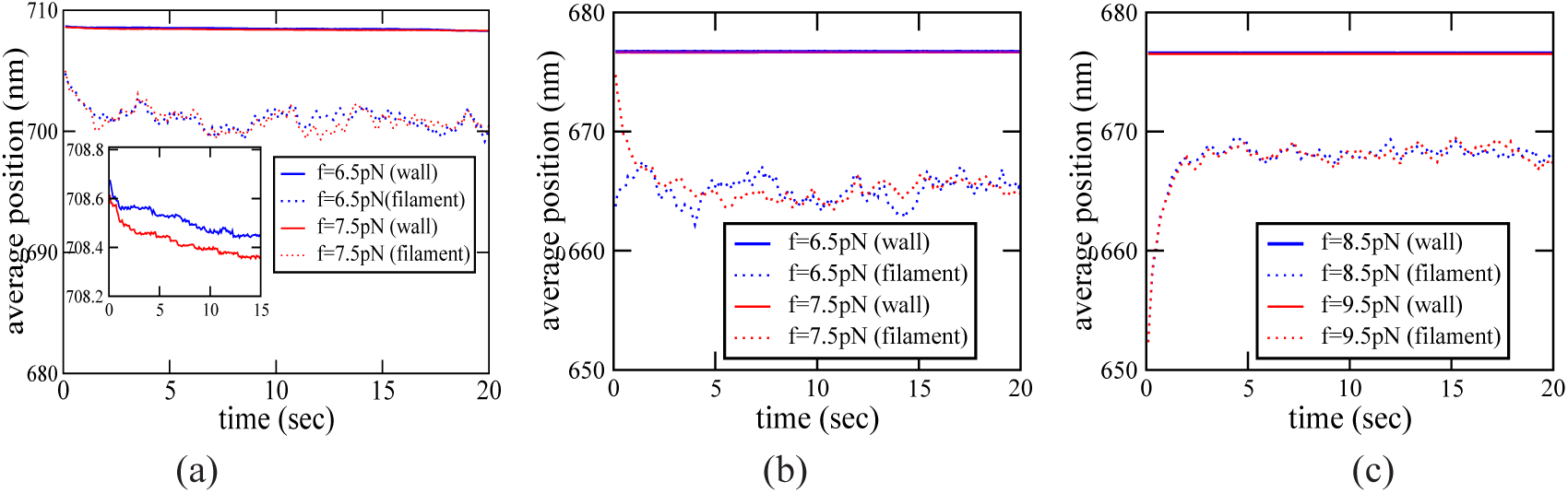
The time evolution of the average positions of the wall and one of the protofilaments is shown for (a) one microtubule, (b) two microtubules (near) and (c) two microtubules (far). In the inset of (a), the average position of wall alone shown. The parameters common for all the three cases are listed in Table 1. For two filaments, ‘near’ means zero base separation, while ‘far’ refers to a base separation 150 nm.

The deviation from linear scaling of the stall force may be characterised using a scaling parameter *ν* = 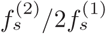, which is always 1 for perfect linear scaling. In Table 2, we collect together the different values of stall forces observed in our simulations, as well as the computed *ν*, for FFD filaments and multi-stranded microtubulelike filaments.

**Table 1:** Numerical values of the various parameters used in the Brownian dynamics simulation for flat-faced filaments (FFD) and microtubule-like geometry.

| Parameter | Symbol | FFD | MT-like geometry |
| --- | --- | --- | --- |
| Monomer length | $\delta$ | 2 nm | 8 nm |
| Radius | $a$ | variable | 2.5 nm |
| Diffusion coefficient(monomer) | $D$ | $10^5 \text{ nm}^2\text{s}^{-1}$ | $10^5 \text{ nm}^2\text{s}^{-1}$ |
| Diffusion coefficient(wall) | $D_w$ | $10^3 \text{ nm}^2\text{s}^{-1}$ | $10^3 \text{ nm}^2\text{s}^{-1}$ |
| Concentration | $C_0$ | $3.3 \mu\text{M}$ | $33.3 \mu\text{M}$ |
| Monomer dissociation rate | $k_{\text{off}}$ | variable | $3 \text{ s}^{-1}$ |

**Table 2:** The stall forces for single and two filaments, and the scaling parameter 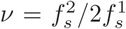, for *N* =1 and 2, for flat-faced disk polymers and microtubule-like polymers. The deviation from unity indicates sublinear scaling with *N*. The error bar in the stall force data is estimated as half of the step size for force used in the simulations. For two filaments, ‘near’ refers to zero base separation and ‘far’ refers to base separation 10 times the radius of cross-section (except for the last, where the base separation is 150 nm).

| Filament | $f_s^1$(pN) | $f_s^2$(pN) | $\nu$ |
| --- | --- | --- | --- |
| FFD ($a = 2$ nm, near) | $2.27 \pm 0.05$ | $4.58 \pm 0.05$ | $1.01 \pm 0.022$ |
| FFD ($a = 2$ nm, far) | $2.27 \pm 0.05$ | $4.53 \pm 0.05$ | $0.99 \pm 0.043$ |
| FFD ($a = 10$ nm, near) | $1.94 \pm 0.05$ | $3.05 \pm 0.05$ | $0.79 \pm 0.026$ |
| FFD ($a = 10$ nm, far) | $1.94 \pm 0.05$ | $3.73 \pm 0.05$ | $0.96 \pm 0.012$ |
| FFD ($a = 20$ nm, near) | $1.88 \pm 0.25$ | $2.76 \pm 0.25$ | $0.73 \pm 0.075$ |
| FFD ($a = 20$ nm, far) | $1.88 \pm 0.25$ | $3.55 \pm 0.25$ | $0.94 \pm 0.059$ |
| Microtubule-like geometry (near) | $5.17 \pm 0.25$ | $6.48 \pm 0.25$ | $0.63 \pm 0.18$ |
| Microtubule-like geometry (far) | $5.17 \pm 0.25$ | $8.02 \pm 0.25$ | $0.76 \pm 0.14$ |

## 4 Discussion

Polymerization-driven force generation by filaments has many biological applications, and the problem has been extensively studied experimentally as well as theoretically. A central quantity of interest here is the stall force of a bundle of *N* filaments, and its scaling behavior with *N*. Recent years have seen a spurt of activity in theoretical modeling in this field, but these models are typically one-dimensional in nature, and do not consider explicitly monomer diffusion in space [18–21, 27], or even the dynamics of the object (barrier) that is being pushed by the filaments [19–21, 27]. Here, we have introduced and studied a more general model in which the effects of polymerization-driven growth of the filament and the presence of the physical barrier on monomer concentration are included, and the consequent fall in the monomer adsorption rate is estimated. We showed that, in general, the physical barrier affects the monomer concentration profile, causing a drop in the growth rate in addition to being a steric hindrance to growth, when the filament tip and the barrier are within a distance of approximately 3-4 times the radius of cross-section of the filament tip (imagined as having a solid disk-like face). We then investigated, by mathematical analysis as well as Brownian dynamics simulations, how the collective dynamics of a bundle of filaments is affected by this barrier-induced hindrance to free diffusion. In the process, we also encountered diffusive interaction between filaments that naturally appears when nearby filaments grow together by diffusion-limited adsorption, but its effects are particularly noticeable in the presence of a barrier as the latter reduces the spread in length across different filaments and thereby forces the tips to be close to each other.

In the mathematical part of our study, we set up a continuum Fokker-Planck equation to describe together the collective growth of *N* identical filaments against a rigid barrier, the latter’s motion including drift towards the filaments and random diffusive motion powered by thermal noise. We then use an adiabatic approximation where the stationary monomer density profile is assumed to respond instantly to changes in the positions of the filament tips and the barrier. The on-rate for adsorption of monomers onto a filament tip is calculated, and also measured directly in simulations. By assuming a simple analytical form for the boundary-affected reduced on-rate, consistent with observations, we studied steady state properties of the filament population. In particular, we derived analytical expressions for the mean filament growth velocity and the stall force of an assembly. These expressions involve the probability distribution for the filament tip-barrier separations (‘gaps’), which was calculated explicitly in a few special cases of interest. All analytical predictions were subjected to verification in Brownian dynamics simulations, which were also used to explore the consequences of having more complex microtubule-like multistranded structure for the filaments.

Among the important conclusions arising out of this study, we have established clearly that, in general, a physical barrier may be expected to cause reduction in the rate of growth of a polymer growing towards it, and this effect also reduces the stall force of the filament. However, as long as the filaments have sufficient lateral separation from each other, the stall force for *N* filaments increases linearly with *N*. Nonlinear scaling appears when the filaments are brought close together to form a bundle; in this case, diffusive interaction between the growing filament tips leads to a non-additive combined stall force of the bundle. The effects of this diffusive interaction are most visible in a multi-stranded filament like a microtubule; here, we show conclusively in simulations that the net stall force of the filament is much less than the sum of the stall forces of the individual protofilaments [6]. Similarly, the combined stall force for two microtubules is generally less than twice the stall force of one. Diffusive coupling, when significant, leads to sublinear scaling of stall force with the number of filaments, a notable prediction from our studies. Specifically, in microtubules, we report the existence of strong diffusive coupling between different protofilaments, arising by virtue of their tight packing, which leads to smaller combined stall force, compared to a hypothetical situation where each protofilament grows independent of the others. We also observe a remarkable phenomenon in our simulations; two microtubules, when growing against a *super-stall* force, stand their ground after retreating to a new ‘equilibrium’ position, and refuse to be continuously pushed backward unlike a single microtubule, or simpler (single-strand) flat-faced filaments. At the moment, we lack a clear understanding of the mechanism or the implications of this observation, and investigating it further is one of our immediate goals for the future.

Among the limitations of our study, we have treated diffusing monomers as point point particles devoid of size and shape; therefore, Brownian rotation of monomers and orientational constraints to their adsorption at the growing tip have been ignored. We do not believe that this will impact our principal conclusions, but if taken into account, could reduce the on-rate uniformly everywhere. Yet another important omission in our model, in the context of microtubules, is that we have not included GTP hydrolysis and the consequent dynamic instability. Recent theoretical work [21] has shown that the combined stall force of a bundle of *N* microtubules with dynamic instability scales superlinearly with *N*. Bundle catastrophes, observed in microtubules growing close together [10] seems to be a collective catastrophe phenomenon which could be studied further using the approach developed in this chapter. In general, it would be interesting to see how the competition between diffusive coupling and dynamic instability, which appear to have opposite effects on the scaling of force with number, affects collective force generation and dynamic instability in a microtubule bundle.

## 5 Authors Contributions

JV and MG contributed equally in designing the project, developing the analytical methods and writing the paper. JV designed and performed the numerical simulations.

## 6 Acknowledgements

We acknowledge P.G. Senapathy Centre for Computing Resources, IIT Madras for computing facilities. We also acknowledge helpful discussions with Dibyendu Das and Ranjith Padinhateeri.

## A Modification of monomer binding rate by the wall

Assuming the filament to be a linear chain of monomers, the probability that a monomer adds in the time interval dt is *k*_on_(*d*)*dt*, where *k*_on_(*d*) is the rate at which monomers are added to the tip. In our model, *k*_on_(*d*) in general depends on the separation between the filament tip and the barrier, denoted as d. When the filament tip is far away from the barrier, *k*_on_(*d*) → *k*_on_ (∞), equivalent to the case studied earlier[18–21]. Due to the steric hindrance arising out of the presence of the wall, a monomer can be added only if there exists a sufficient space between the filament tip and the barrier, equal to the size of the monomer.

Assuming steady state conditions, the monomer concentration *C*(**r**, *t*) satisfies Laplace’s equation

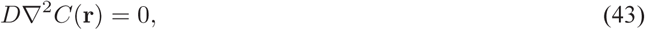
where **r** is the position measured with respect to the centre of the surface and *D* is the diffusion coefficient of monomers. In diffusion-limited growth, the steady state adsorption rate of monomers to a surface *S* is given by the integral

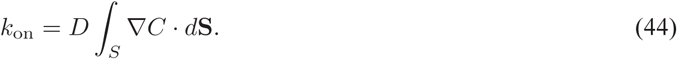

To find *C*(**r**), we solved Eq.43 in a geometry (having cylindrical symmetry) as shown in Fig.9a, consisting of an absorbing disk of radius *a* and zero thickness placed at *z* = 0 and an infinite reflecting boundary at *z* = *d*. Given this geometry, the solution of Eq.43, denoted as *C_d_* (**r**), is given by

**Figure 9:**
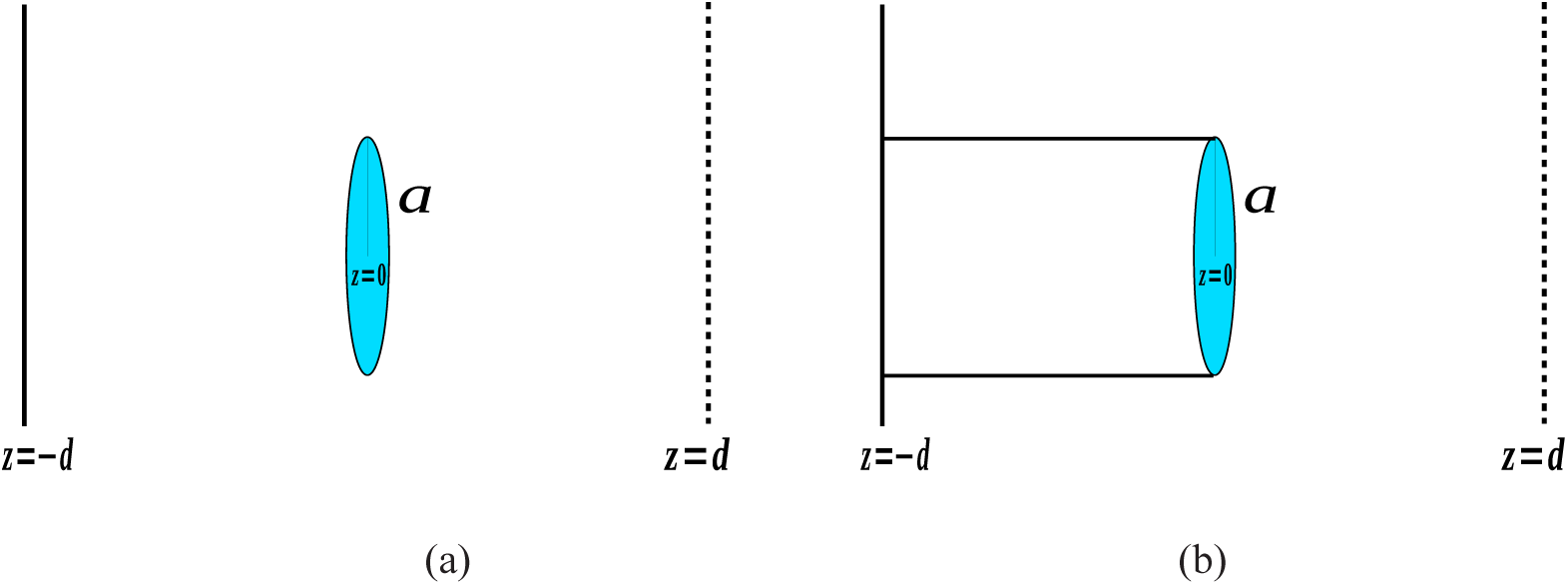
(a)Schematic figure of the geometry used to solve Eq.43. A thin circular disk of radius *‘a’* kept in between two rigid infinite walls, which acts as an absorbing region for the incoming particles. (b) Schematic figure of the geometry used in Brownian dynamic simulations. In addition to the conditions in (a), a reflecting boundary condition is imposed on the cylindrical wall.

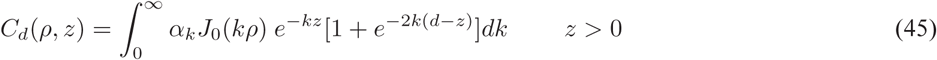

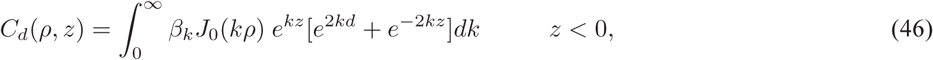
where *α_k_* and *β*_k_ are constants to be fixed and the subscript *d* indicates the location of the barrier. To see how the presence of the reflecting wall at *z* = *d* affects the on-rate of particles coming from *z* > 0, we evaluate the integral in Eq.44 using Eq.45. The constant *α_k_* in Eq.45 is fixed such that *C_d_*(*ρ, z*) in the positive *z* region satisfies the full set of boundary conditions:

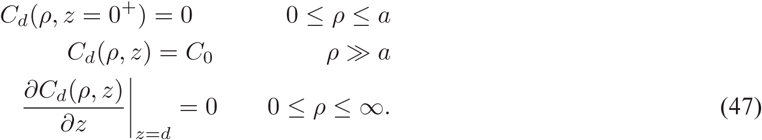

Consistent with the above boundary conditions, the solution in the region *z* > 0 becomes,

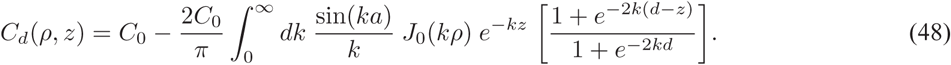

As a special case, for *z* = *d*, the solution, obtained after performing the integration in Eq.48 is

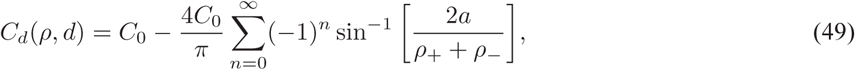
where

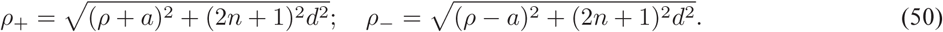

For comparison, if the wall were not present, the corresponding solution (again, at *z* = *d*) would be

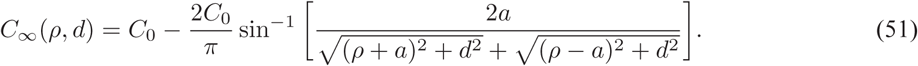

In Fig.10, we show the concentration profile (dimensionless, scaled using the asymptotic value *C_0_*) in the radial direction, given by Eq.49 and Eq.51, for *a* = 20 nm and *d* = 10 nm. The figure shows that the presence of the wall enhances monomer depletion in front of the growing filament tip, and this effect is found over a (radial) distance nearly 4 times the radius of cross-section of the absorbing disk. It is natural to expect that this depletion will also cause a fall in the rate of adsorption of the monomers at the disk, which we calculate next using Eq.44.

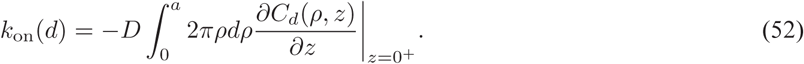

**Figure 10:**
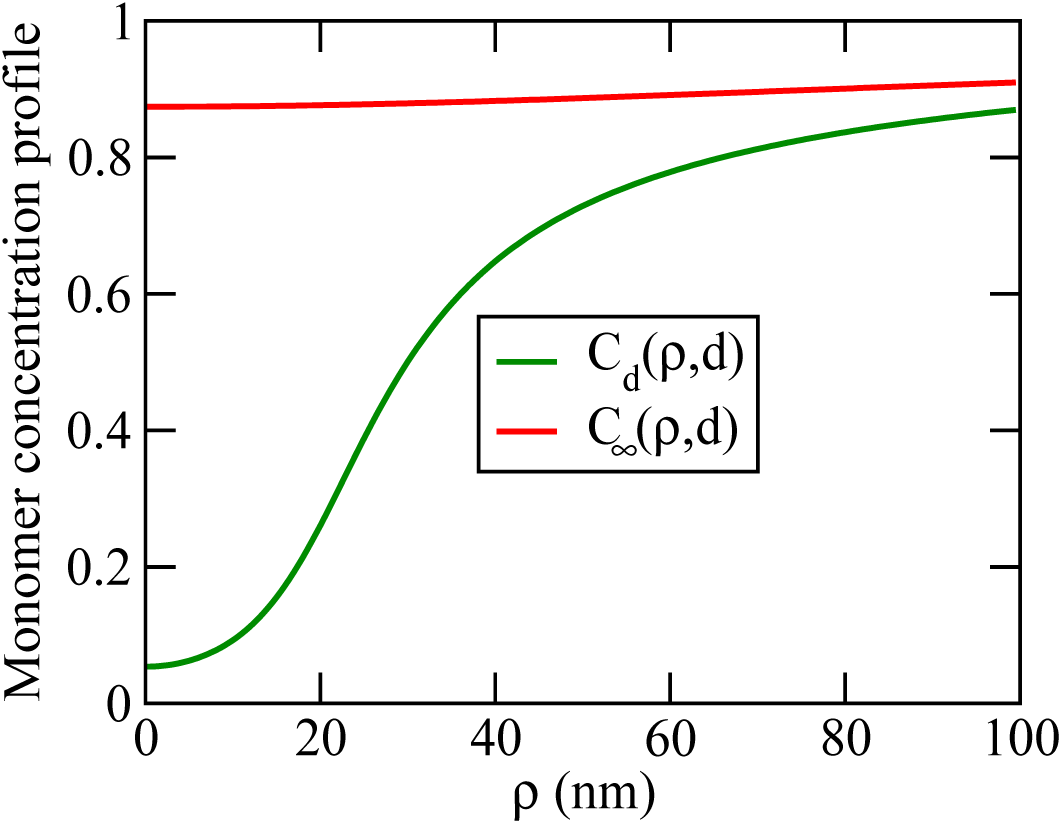
A comparison of the concentration profile of free monomers along the radial direction, given by Eq.49, and Eq.51, for disk radius *a* = 20 nm at *z* = 10 nm, when the reflecting barrier is placed at *z* = 10 nm (green) and *z* = ∞(red). Far away from the barrier, the concentration is given by the asymptotic value *C_0_*. Note that the presence of the barrier enhances depletion of monomers in front of the absorbing disk.

Using the expression for *C_d_* (*ρ, z*) in the region *z* > 0 given by Eq.48, we have

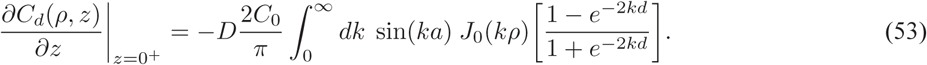

Substituting Eq.53 in Eq.52 and performing the integration, we find

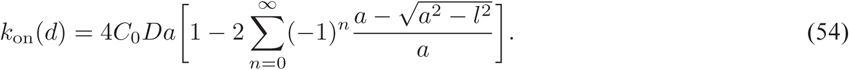
with

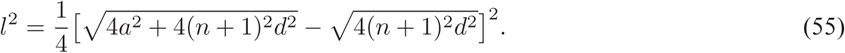

In the limit *d* → ∞ (disk far away from the barrier), the quantity *l*^2^ given by Eq.55 goes to zero and the on-rate takes the simple expression [28],

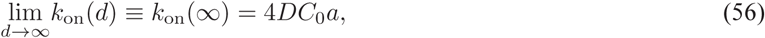
as expected.

The on-rate decays monotonically as the wall-disk separation decreases, suggesting that the presence of a barrier decreases the likelihood of particles being getting trapped and hence slows down the growth rate, see Fig. 11. We also verified the prediction for on-rate given by Eq.54, by doing Brownian dynamics simulations, for more details see section on simulations case (i) discussed in Sec.2.3. A visual inspection of Fig.11 suggests that boundary induced drop in on-rate comes into play when the separation between the barrier and radius of cross section of the absorbing disk are comparable, i.e., *d ∼ a*. Unfortunately, the expression for the flux given in Eq.54 is not simple enough to be used directly for further mathematical calculations, hence we approximate Eq.54 by the simpler form: *k*_on_ ∼ *k*_on_ (∞)(1 – *e*^−λ*d*^). The inverse of the parameter λ gives a measure of the size of depletion zone, i.e., λ ∼ 1/*a*. In Fig.11b, we give a fit of the approximate expression for on-rate with the simulation data for *a* =10 nm. The best fit parameter value of λ in this case is 0.275 nm^−1^.

**Figure 11:**
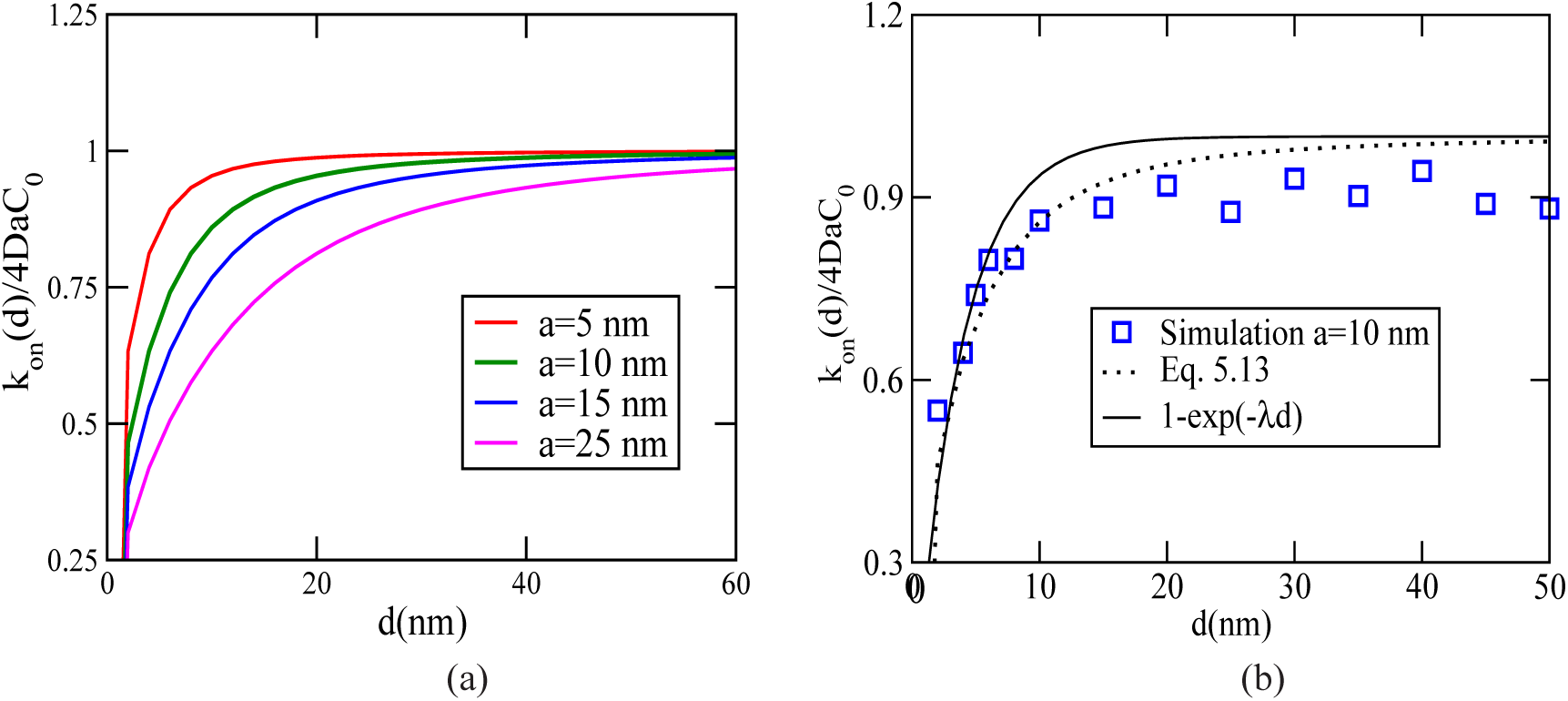
(a) On-rate of particles to a circular disk of radius ‘*a*’ as a function of distance between the disk and the reflecting wall for different disk radii, given by Eq.54. (b) A comparison of analytical result for the on-rate given by Eq.54 with results from Brownian dynamics simulations discussed in Sec.3.3. The Brownian dynamics simulations are carried out for a geometry shown in Fig.9b; for more discussions on details of simulations, see Sec. 2.3. The thick line is shown for the approximated scaled expression of the on-rate, *k*_on_ (*d*)/*k*_on(*∞*)_ = 1 – exp(–λ*d*) with the best fit value of λ = 0.275 nm^−1^. The dotted line is the analytical expression given by Eq.54.

## B 1/λ expansion for *V_N_* (*f*) and 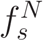

As a simple extension of the calculation discussed in Sec.3.2, an asymptotic expansion of the expressions for average filament velocity and stall force in the large λ limit is carried out, to quantify small deviations from the constant on-rate case. Consider the equation for single filament gap distribution, *ϕ_N_*(*y*) given by Eq.20. Performing an integration with respect to *y* in Eq.20, and using Eq.S15 and Eq.S16 we get,

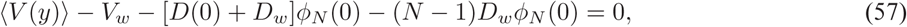
which yields

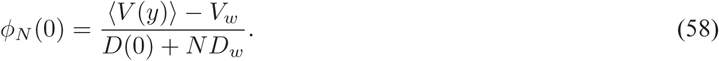

Substituting Eq.58 in the expression for *V_N_* (*f*) given by Eq.13 gives

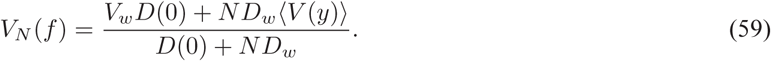

In general, *ϕ_N_*(*y*) is a function of λ as well, therefore let us express *ϕ_N_*(*y*) explicitly as a function of λ and *N* i.e.,

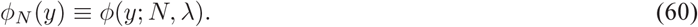

The quantity 〈*V*(*y*)〉 is calculated using Eq.25,

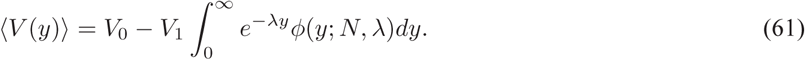

Next, we expand *ϕ*(*y*; *N*, λ) in powers of λ:

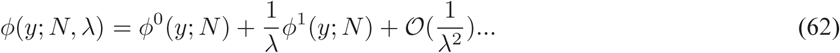

From Eq.38, we have

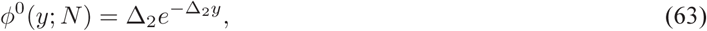
where Δ_2_ is given by Eq.35. To evaluate 〈*V*(*y*)〉, we substitute Eq.62 in the integral given in Eq.61, also substitute *ϕ*^0^(*y*; *N*) from equation Eq.63, and we find

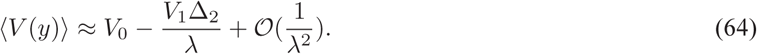

Therefore, from Eq.59, we have

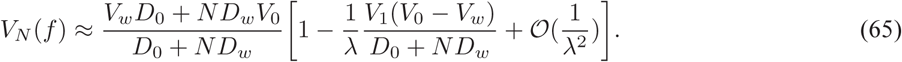

The stall force is obtained from Eq.65 using the defining relation 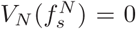, and takes the form of an powe-series expanision in 1/λ:

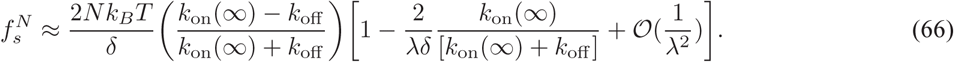

From Eq.65 and Eq.66, it is evident that the barrier-induced inhibition of free diffusion causes a drop in the mean velocity and stall force, while the linear scaling of stall force with the number of filaments holds, at least to first order in 1/λ. Numerical simulations, discussed in the main text, indicate that this result holds for arbitrary λ, under conditions where the filaments grow independent of each other.

